# A pleiotropic flowering time QTL exhibits gene-by-environmental interaction for fitness in a perennial grass

**DOI:** 10.1101/2022.02.26.482116

**Authors:** Xiaoyu Weng, Taslima Haque, Li Zhang, Samsad Razzaque, John T Lovell, Juan Diego Palacio-Mejía, Perla Duberney, John Lloyd-Reilley, Jason Bonnette, Thomas E Juenger

## Abstract

Flowering time is crucial for wild plant populations to adapt to their local environments. Although the genetic basis of flowering variation has been studied in many plant species, its mechanisms in non-model organisms and its adaptive value in the field are still poorly understood. Here, we report new insights into the genetic basis of flowering time and its effect on fitness in *Panicum hallii*, a native perennial grass. We conducted genetic mapping in populations derived from representative inland and coastal ecotypes to identify flowering time QTL and loci exhibited extensive QTL-by-environment interactions. Patterns of segregation within recombinant hybrids provide strong support for directional selection driving ecotypic divergence in flowering time. A major QTL on chromosome 5 (*q-FT5*) was detected in all experiments and is a key locus controlling flowering variation. Fine-mapping and expression studies identified a *FLOWERING LOCUS T* orthologue, *FT-like 9* (*PhFTL9*), as the candidate underlying *q-FT5*. We used reciprocal transplant experiment to test for global local adaptation and the specific impact of *q-FT5* on performance. We did not observe local adaptation in terms of fitness tradeoffs when contrasting ecotypes in home versus away habitats. However, we observed that the coastal allele of *q-FT5* conferred a fitness advantage only in its local habitat but not at the inland site. Sequence analysis of the *PhFTL9* promoter identified ecotypic specific *cis*-element variation associated with environmental responsiveness. Together, our findings demonstrate the genetic basis of flowering variation in a perennial grass and provide evidence for conditional neutrality underlying flowering divergence.

## Introduction

Local adaptation is a key component of responses to changing environments and a central topic in modern evolutionary biology (1). Plants provide unique opportunities to study the ecological and evolutionary history of local adaptation, as they are unable to escape from danger and must tackle the challenges that local conditions present (2). The selection of favored phenotypes by local habitats usually leads to phenotypic differentiation among natural populations, thereby enhancing fitness and promoting local adaptation (3-5). One common hypothesis is that local adaptation driven by natural selection results in trade-offs involved in specialization (6, 7). Alleles increasing fitness in one environment may through antagonistic pleiotropy result in decreases in fitness in other environments. Alternatively, local adaptation may arise through mutations that improve fitness locally, but that are neutral and generally have no fitness impact in other environments (i.e. conditional neutrality) (8-10). Determining the frequency of antagonistic pleiotropic versus conditionally neutral allelic effects is important for understanding the forces that maintain standing genetic variation, the importance of gene flow and recombination in facilitating or constraining adaptation, and the likelihood or extent of local adaptation in natural populations (11). While progress has been made (12), too few empirical studies have explored the genetic architecture of natural alleles and their interaction with native environments. Therefore, field studies bringing together the genetic basis and ecological significance of ecologically important traits are critically needed.

Timing the reproductive transition is a key developmental decision in the life history for all species (13-15). In plants, depending on geography, the patterns of flowering time generally evolve in response to local environmental factors like the day-length, temperature, and different types of biotic or abiotic stresses through changes in flowering time pathway genes (16-18). As a complex trait, flowering time is often tightly integrated within the genetic networks of other traits related to plant growth and stress responses (19). Although it is not always clear whether this phenomenon is due to selection on flowering time itself or through other traits sharing the same genetic network, pleiotropy (one gene affecting multiple traits) is one of the most commonly observed attributes of genes involving flowering time pathways (19). For example, several main flowering time genes in *A. thaliana*, including *FT* and its regulators (e.g. *FRIGIDA* (*FRI*) and *FLOWERING LOCUS C* (*FLC*)), have shown pleiotropic effects on development (e.g. branching architecture and seed germination) and physiological (e.g. water use efficiency) characteristics (20-22). In rice, several *FT* homologs (*HEADING DATE 3a* (*Hd3a*) and *RICE FLOWERING LOCUS T 1* (*RFT1*)) and their upstream regulators (*HEADING DATE 1* (*Hd1*) and *GRAIN NUMBER, PLANT HEIGHT AND HEADING DATE 7* (*Ghd7*)) have roles in vegetative and reproductive branching development (23-26). These results suggest potential adaptive roles of flowering time genes in the maintenance of diverse life-history strategies across different populations.

Determining the adaptive value at the genetic loci or even the gene level is critical to understand the molecular basis of local adaptation (27). To date, extensive inquiry into the genetics of adaptation have revealed quantitative trait loci (QTL) in many systems (9, 28, 29). These studies are especially potent in genetic models, such as *A. thaliana*, where the putative function of many genes can be well characterized (28, 29). However, most such studies never test the adaptive value of individual genetic loci in the environments in which the traits evolved. In terms of the gene level, although more than 300 genes involved in flowering time regulation have been identified in *A. thaliana* (30), only a few of them have been tested for fitness effects with mutants grown under realistic natural conditions (31, 32). For non-model species, unfortunately, the challenge is more apparent because of the lack of genetic resources (e.g. mutants or genetic population) and related knowledge of the genetic architecture of target traits (e.g. QTL or candidate genes).

*Panicum hallii* (*P. hallii*) is a genetically tractable native perennial grass occurring in North American with a geographic range that spans a number of ecoregions and native habitats (33). Two major ecotypes of *P. hallii* are found in inland (*var. hallii*) or coastal (*var. filipes*) habitats across the southwest (34, 35). Consistent with differential adaptation to xeric inland and mesic coastal habitats, the HAL2 genotype, which is representative of *var. hallii* ecotype, flowers earlier and develops faster than the FIL2 genotype, representing the var. *filipes* ecotype (34). Independent *de novo* genome assemblies and annotations have been built for the HAL2 and FIL2 accessions and F_2_ and recombinant inbred line mapping populations have been generated between these two genotypes (34, 36, 37). In this study, we leverage the resources of *P. hallii* to investigate the genetic basis of flowering time and test the adaptive value of a major genetic factor under field conditions. We specifically ask the following four questions: whether flowering time has evolved under natural selection in *P. hallii*? What genomic regions contribute to flowering time divergence and genotype-by-environment (G x E) interaction between representative inland and coastal ecotypes? Does the major flowering time QTL impact fitness and responsible display trade-offs or conditional neutrality under natural field conditions? Do certain mutation types (e.g. *cis*-regulatory changes) preferentially contribute to flowering time adaptation on the major genetic factor?

## Results

### Genotype-by-environment interactions and natural selection for flowering time

To probe the genetic architecture of evolved differences in flowering time between the ecotypes, we applied quantitative trait locus (QTL) mapping across different seasons using an F_2_ and recombinant inbred line (RIL) populations derived from HAL2 and FIL2. Despite substantial variation of photoperiod among the four controlled experiments in which flowering time was assayed, the HAL2 parent always flowered significantly earlier than the FIL2 parent (t-tests, *P* < 0.01 from all four experiments) (Fig. 1 and Table S1). In the F_2_ population, the difference in mean flowering time between parents was 25.8 days (Cohen’s ds = 8.3), and this difference in three RIL populations ranged from 8.0-25.3 days (Cohen’s ds ranged from 3.2 to 7.4) (Table S1). The broad-sense heritability (*H*^*2*^) of flowering time in three RIL experiments in 2015 fall, 2016 spring, and 2016 summer were 0.42, 0.31, and 0.41, respectively. We observed positive genetic correlations less than a value of 1 across three seasonal RIL experiments (*r*_*g-fall-spring*_ = 0.56, *r*_*g-fall-summer*_ = 0.65, *r*_*g-spring-summer*_ = 0.60), suggesting the presence of genotype-by-environment (G x E) at the trait level. We further tested for G x E across the RIL experiments using factorial linear models and detected a strong G x E interaction (*P* < 0.001) of flowering time in three seasons of study. These results show that flowering time variation between representatives of each ecotype was heritable and sensitive to different seasonal environments, potentially resulting from differential environmental sensitivity at discrete genetic loci. We used the Fraser *v*-test statistic based on segregating genetic variance to test for directional selection as an explanation for flowering time divergence. In all cases, the variances of parental flowering time values were greater than the variances among RILs and *v*-tests were highly significant (in all cases, *P* < 0.05) for flowering time measured in four QTL experiments (Table S2). These results suggest historical directional selection underlying flowering time divergence among *P. hallii* ecotypes.

**Fig. 1.**
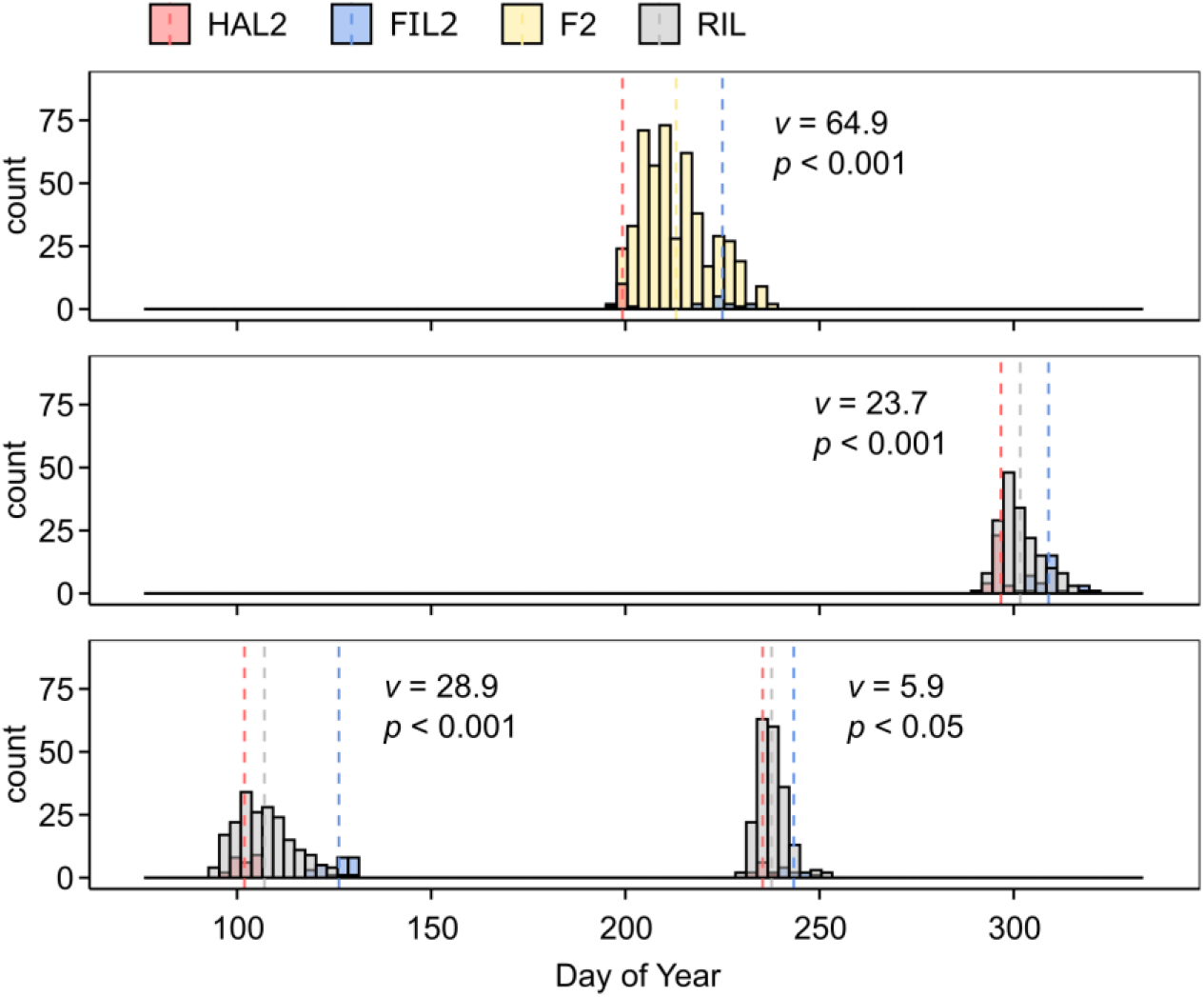
Flowering time variation of *P. hallii* in F_2_ and RIL populations. The distribution of flowering time in F_2_ population in 2014 (above), and RIL population in 2015 (middle) and 2016 (bottom) experiments. The parental HAL2 (triangle) and FIL2 (cross) means are plotted above each distribution.

### Mapping QTL for flowering time in an F_2_ population

To investigate the genetic basis of flowering divergence between the ecotypes, we sequenced 2 bulk populations with extremely early or late flowering times (EF-pool and LF-pool), each consisting of ∼20% of the progeny from the tail of the F_2_ population. After mapping and SNP calling, we explored allele frequency shifts in the bulked progeny by computed a SNP-index for the EF and LF pools as well as their differences, Δ (SNP-index). Two regions on chromosome 3 and 5 had an average SNP-index higher than 0.7 in EF-pool and one region on chromosome 5 had an average SNP-index as low as 0.25 in LF-pool (Fig. 2). By examining the Δ (SNP-index) plot, we identified two genomic regions exhibiting the highest Δ (SNP-index) values: the region on chromosome 3 from 0.3 to 9.9 Mb and the region on chromosome 5 from 0.5 to 11.1 Mb (95% confidence intervals) (Fig 2). To further confirm the flowering QTL detected by QTL-seq, we developed 13 insertion/deletion (InDel) markers in two regions (Table S3) and conducted a classical bi-parental QTL analysis within the same F_2_ population. We found the peak LOD scores at the marker M3-4979 on chromosome 3 (LOD = 15.3) and the marker M5-8935 on chromosome 5 (LOD = 12.1). Our results in the F_2_ population suggested that flowering time is controlled by a few major-effect QTL on chromosome 3 and 5 in *P. hallii*.

**Fig. 2.**
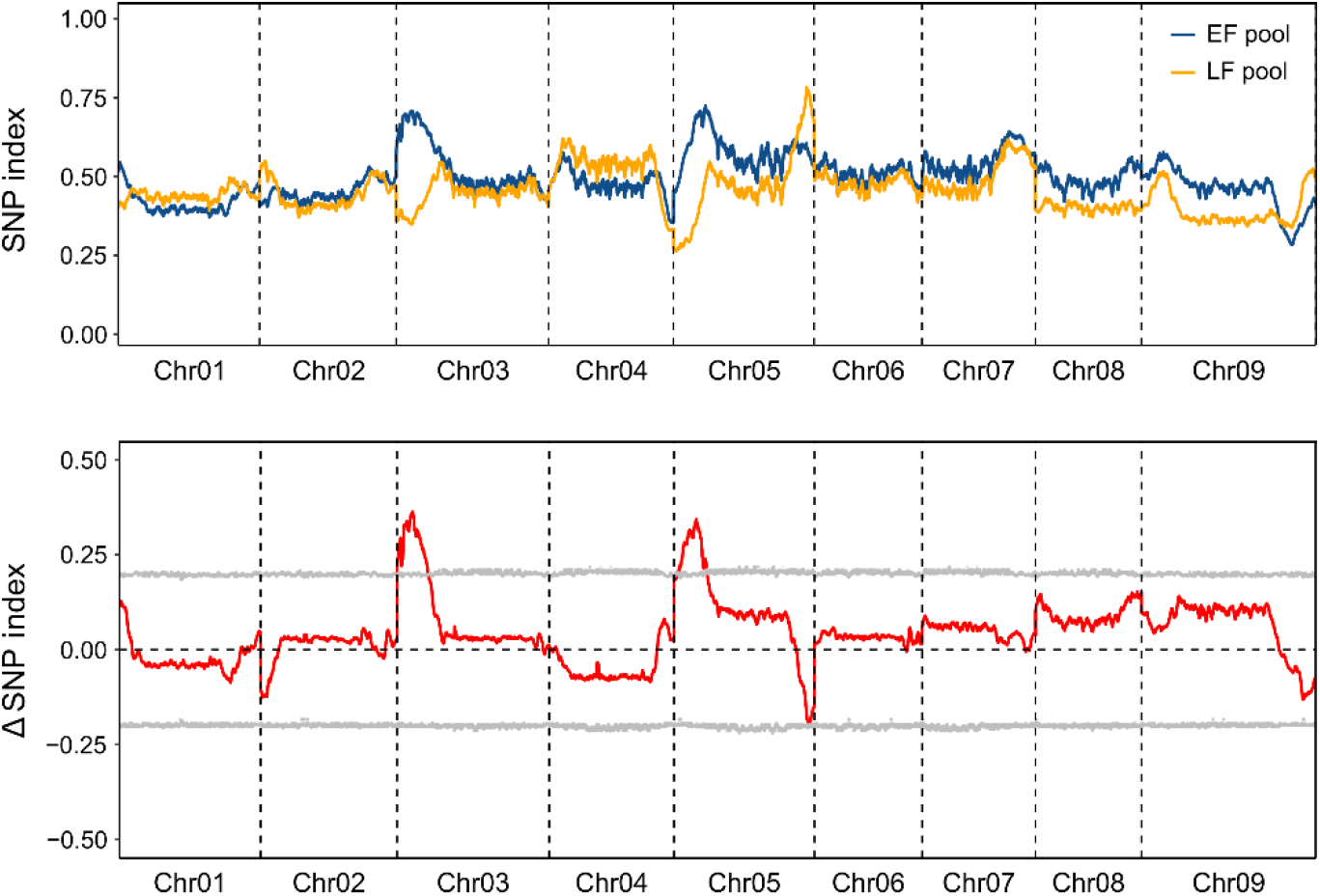
SNP index plots of early flowering (EF) and late flowering (LF) population and Δ (SNP index) plots generated by sliding-window analysis. The SNP indexs (ratio of the SNPs that are identical to those in the EF and LF pools) are shown using blue and orange lines, respectively. The Δ (SNP index) (subtracting the SNP index of the LF population from that of the EF population) and its 95% confidence interval are shown using red and grey lines, respectively.

### Mapping QTL for flowering time in a RIL Population

To dissect genomic regions governing flowering differences and patterns of QTL-by-environment interaction (QTL x E), we mapped QTL in the RIL population grown across three experimental conditions using a modeling strategy incorporating QTL x E. QTL x E was tested as the difference between a full model incorporating seasonal environments as an interactive covariate versus a reduced model only controlling for additive effects of the environments. These analyses confirmed the major QTL on chromosomes 3 and 5 along with three novel QTL detected in the full model. (Fig. 3 and Table S4). Further, by comparing the full model and the reduced model, we identified nine genomic regions exhibiting significant QTL x E, including all QTL detecting in the full model and four other regions on chromosomes 1, 2, 4, and 6 (Fig. 3 and Table S4). The QTL effects showed that HAL alleles of all five QTL in the full model accelerated flowering in most seasons, except the QTL on the bottom of chromosome 3 in 2015 fall and 2016 summer (Fig. S1). Moreover, we observed stronger QTL effects for all five QTL in 2016 spring relative to the two other seasons (Fig. S1).

**Fig. 3.**
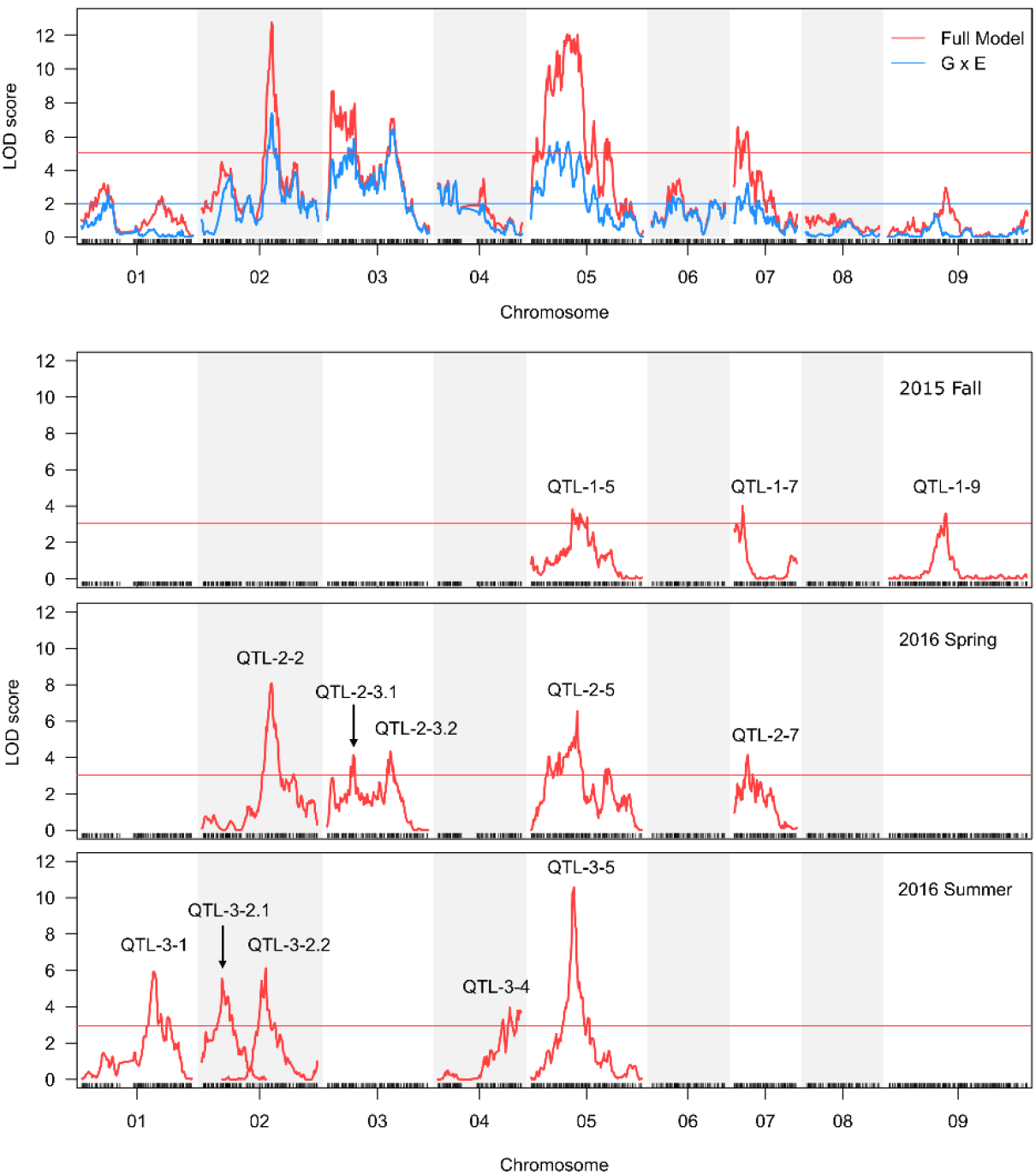
Location of flowering time QTLs under different environmental conditions. QTL x E was tested as the difference between full model including QTL x E interaction and a reduced additive model (Above). The red lines represent the QTL identified in the full model. The blue lines represent the QTL x E threshold identified by model comparisons. QTL mapping for each RIL experiment are shown as below. The red horizontal line represents the threshold of significance based on permutations. The name attributed to the different QTLs are shown above the QTL peak.

To further explore the genetic basis of flowering variation in each seasonal condition, we performed multiple QTL mapping based on stepwise modeling for each experiment separately. Here, we notate QTL by their season of occurrence and their chromosome location (QTL-season-chromosome). We detected three QTL in 2015 fall (QTL-1-5, QTL-1-7, and QTL-1-9), five QTL in 2016 spring (QTL-2-2, QTL-2-3.1, QTL-2-3.2, QTL-2-5, and QTL-2-7), and five QTL in 2016 summer (QTL-3-1, QTL-3-2.1, QTL-3-2.2, QTL-3-4, and QTL-3-5) (Fig. 3 and Table S5). The majority of QTL had additive effects with FIL2 alleles delayed flowering, except two QTL (QTL-3-1 and QTL-3-2.1) detected in 2016 summer (Table S5). The proportion of total variation in flowering time explained by significant QTL in each experiment varied from 24.3% to 43.5% (Table S5). The additive effects of each QTL were 0.78-2.73 days and explained 5.9-17.8% of the phenotypic variation (Table S5). Among these, the QTL on chromosome 5 was identified across three seasons of study (QTL-1-5, QTL-2-5, and QTL-3-5) with the highest peak LOD scores during the summer of 2016 (QTL-3-5, LOD = 10.6) (Fig. 3 and Table S5). Interestingly, we detected an epistatic interaction between QTL-3-5 and QTL-3-1 in 2016 summer (Fig. S2). Moreover, we detected three minor QTL (QTL-1-9, QTL-3-1, and QTL-3-4) that were not observed in our QTL x E analysis (Fig. 3 and Table S5). These findings enhance our understanding of the genetic basis of flowering time and highlight the locus on chromosome 5 as a major flowering QTL in *P. hallii*.

### Fine mapping of the major flowering time QTL on chromosome 5

Given the importance of the QTL on chromosome 5 (*qFT-5*), we used a heterogeneous inbred family (HIF) strategy for fine-mapping (38). A RIL line (FH-312) that was heterozygous in the *qFT-5* region was chosen to develop our HIF (Fig. S3). First, we observed that the HIF progenies carrying FIL2 homozygous alleles at *qFT-5* region displayed a significantly later flowering time than those carrying HAL2 homozygous alleles (t-test, *P* < 0.01; Fig. S4), suggesting that the causal region of *qFT-5* is located between two flanking markers. To refine the physical interval, we identified 12 recombinants among the progeny using two flanking markers M5-7948 and M5-12422 (Fig. 4A) and conducted QTL mapping in a recombinant-derived F_3_ population. We detected significant flowering time differences from marker M5-9163 to M5-10558 with the peak at marker M5-9635 (*F* = 27.775, *P* < 0.001; lsmeans-FIL2 = 48.1 ± 0.4; lsmeans-HAL2 = 45.2 ± 0.4; contrast estimate FIL2-HAL2 = 2.9 ± 0.5, *P* < 0.001) (Fig. 4B). It is worth noting that the physical position of marker M5-9635 (9,038,694 - 9,039,322 bp) is only ∼ 30-kb away from the QTL LOD peak (9,068,928 bp) identified in RIL mapping. Accordingly, we created a pair of nearly-isogenic lines (NIL) (NIL^qFT-5-FIL2^ and NIL^qFT-5-HAL2^) spanning a 380-kb region around the peak region. We detected a significant flowering time difference between NIL^qFT-5-FIL2^ and NIL^qFT-5-HAL2^ plants (lsmeans-NIL^qFT-5-FIL2^ = 49.6 ± 0.4; lsmeans-NIL^qFT-5-HAL2^ = 46.3 ± 0.5; contrast estimate NIL^qFT- 5-FIL2^ - NIL^qFT-5-HAL2^ = 3.3 ± 0.7, *P* < 0.001) (Fig. 4C), suggesting that the 380-kb region may harbor the candidate gene for *qFT-5*.We identified 60 and 62 annotated genes in HAL2 and FIL2 genome assemblies, respectively (Table S6). There is no function annotation for two presence/absence genes, and the Ka/Ks estimation suggested the purifying selections for genes in the QTL interval (Table S6). We examined the expression variation of candidate genes in the interval between HAL2 and FIL2 from an existing high replicated transcriptome dataset. This analysis led us to identify 16 differential expression genes, including a phosphatidylethanolamine-binding protein homologous to the rice *FT-like 9* gene (*PhFTL9*) (Table S6). We further performed the qPCR analysis and confirmed that the expression of *PhFTL9* in HAL2 plants was significantly higher than that in the FIL2 plants (Fig. S5). In plants, the members of the *FT* and *FT-like* gene family are major components of florigen and their gene expression variability has been shown to impact flowering time (39-41). Therefore, we suggested that *PhFTL9* is a likely candidate gene underlying *q-FT5*.

**Fig. 4.**
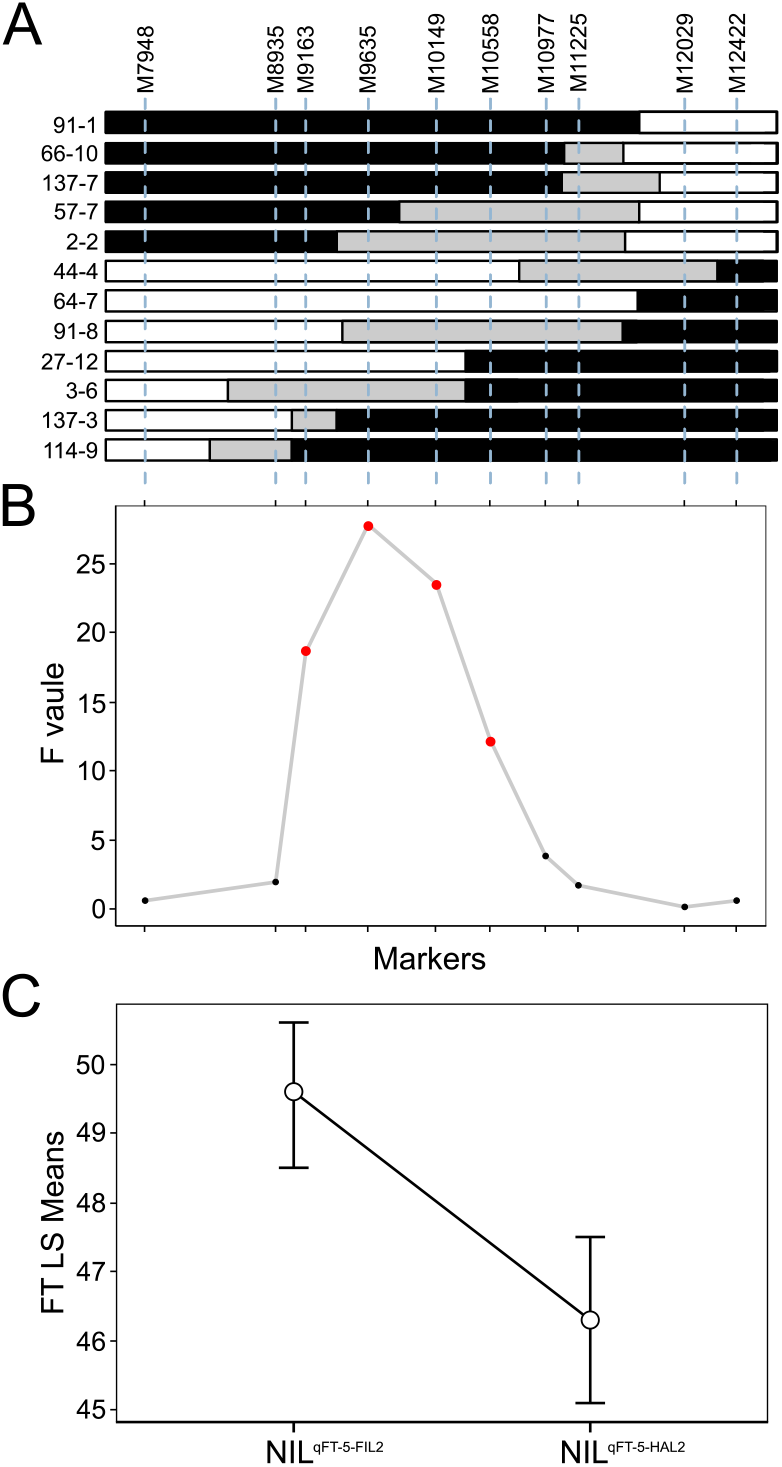
Fine-mapping of the major flowering time QTL, *qFT-5*. A. The recombinants identified from HIF experiments. Black, white and gray boxes indicate homozygous regions for HAL2, homozygous regions for FIL2, and heterozygous regions where recombination occurred, respectively. B. The results of ANOVA examining the effect of genotypes across eight interval markers in recombinant-derived F_3_ families for flowering time. C. The flowering time difference between NIL^qFT-5-FIL2^ and NIL^qFT-5-HAL2^ plants.

### The *qFT-5* conditionally affects fitness-based adaptation in native fields

To investigate the ecological and adaptive significance of *qFT-5*, we performed a field reciprocal transplant experiment using NIL pairs (NIL^qFT-5-FIL2^ and NIL^qFT-5-HAL2^) and two parents (FIL2 and HAL2) grown at their location of collection, Kingsville and Austin Texas, respectively. We measured performance traits related to growth and fitness over the course of a single summer growing season. For both parents, all fitness-related traits were significantly higher at the coastal site than at the inland environment (Fig. 5). The location x genotype interaction for all fitness-related traits was statistically significant in the comparison of parental transplants with a two-way ANOVA test, especially for above-ground biomass per plant (*F* = 357.5, *P* < 0.001) and number of seeds per plant (*F* = 110.1, *P* < 0.001) (Fig. 5 and Table S7). FIL2 outperformed the HAL2 parent for biomass and fitness at both locations, but to an increased degree at the coastal site. This pattern is in conflict with a hypothesis of local adaptation over this summer study season. For NIL pairs, we observed a significant location x genotype interaction for most fitness-related traits except the number of primary and secondary branches per panicle (Table S7). In comparisons between NIL pairs, NIL^qFT-5-FIL2^ plants produced more tiller, biomass, and seeds relative to NIL^qFT- 5-HAL2^ plants at Kingsville (number of tillers per plant mean, NIL^qFT-5-FIL2^ = 121.1 ± 3.6, NIL^qFT-5-HAL2^ = 100.3 ± 2.9, *P* < 0.001; number of secondary branches per panicle mean, NIL^qFT-5-FIL2^ =33.7 ± 0.7, NIL^qFT-5-HAL2^ = 31.4 ± 0.6, *P* < 0.01; number of flowers per panicle mean, NIL^qFT-5-FIL2^ = 397.1 ± 8.2, NIL^qFT-5-HAL2^ = 357.9 ± 7.7, *P* < 0.001; above ground biomass per plant mean, NIL^qFT- 5-FIL2^ = 99.4 ± 4.6, NIL^qFT-5-HAL2^ = 86.6 ± 3.5, *P* < 0.05; number of seeds per plant mean, square root transformation, NIL^qFT-5-FIL2^ = 218.1 ± 5.0, NIL^qFT-5-HAL2^ = 188.7 ± 4.2, *P* < 0.001) (Fig. 5), suggesting a fitness advantage of the local allele at *qFT-5* at the coastal environment during our season of study. In contrast, no statistically significant differences were recorded in fitness-related traits at the inland environment, Austin (Fig. 5). These results suggested that the major flowering time QTL, *qFT-5*, conditionally controls fitness-based adaptation in *P. hallii*.

**Fig. 5.**
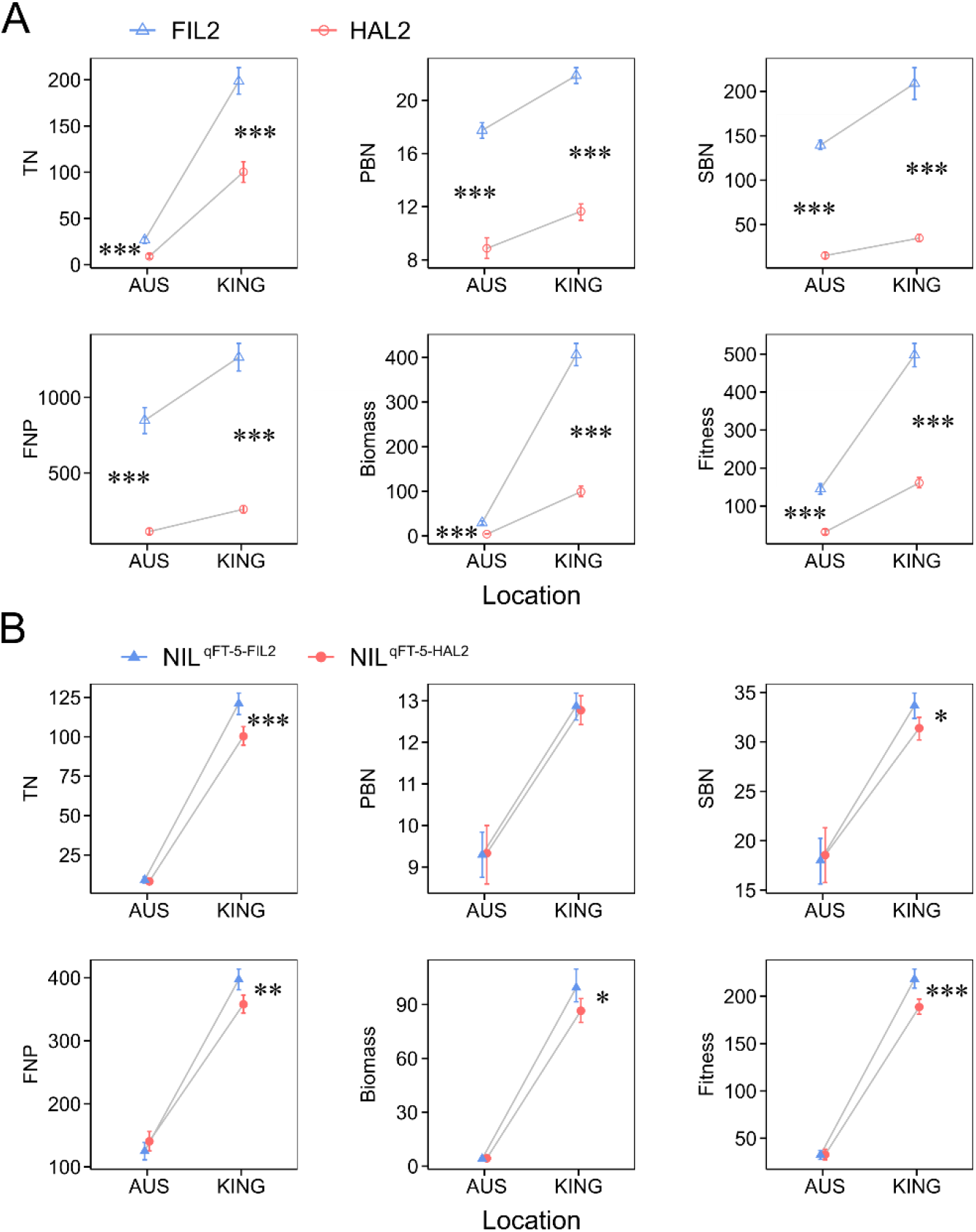
Fitness-related traits of parents (A) and HIF plants (B) in reciprocal transplant experiments. The results of two-way ANOVA examining the effect of location and genotype; contrasts were used to test the effect of genotype separately by site. *, *P* < 0.05; **, *P* < 0.01; ***, *P* < 0.001.

### Sequence variation and evolutionary analysis of *PhFTL9*

To investigate the molecular basis of divergence in *PhFTL9*, we analyzed sequence diversity in the coding and promoter region of the candidate gene. First, we evaluated patterns of synonymous and non-synonymous mutations in the *PhFTL9* protein based on the HAL2 and FIL2 reference genome assemblies. This analysis revealed only two synonymous changes and a single amino acid substitution (S2L) between HAL2 and FIL2 genotypes (Fig. S6). This amino acid substitution is not involved in the known functional regions of the protein, like the potential ligand-binding pocket or the external loop domain (42), suggesting that the function of *PhFTL9* is highly conserved between the two genotypes. Ka/Ks ratio tests of neutrality based on a number of related Panicoid grasses showed that all orthologous pairs had a ratio varying from 0.066 to 0.457 (*P* < 0.01) (Table S8), suggesting that *FTL9* gene has generally undergone purifying selection. Next, we investigated the nucleotide polymorphisms occurring over a 2-kb region of the *PhFTL9* promoter and the 5’ UTR region. We identified a high degree of polymorphism that differentiated FIL2 and HAL2 alleles (Fig. S7). Interestingly, we found a 26-bp deletion at -1265 bp in the HAL2 promoter relative to the FIL2 sequence, which resulted in the loss of a number of motifs involved in light signaling (e.g. SORLIP1) and stress responses (e.g. W-box and core motif of dehydration-responsive element) (Fig. 6A) (43-45). To expand our analyses beyond the parents to a broader sample of both ecotypes, we obtained the *PhFTL9* promoter sequences for 12 *P. hallii* accessions, including four coastal and eight inland ecotypes spanning a wider geographic range of the southwest (Fig. 6B). This Indel is a fixed difference between the inland and coastal ecotypes with a clear geographic differentiation between coastal and inland habitats (Fig. 6B). Previous studies reported that Indel polymorphisms occurring in the *FT* promoter play a critical role in gene expression regulation and ecotype divergence in *Arabidopsis* (41). Therefore, the InDel polymorphism in the *PhFTL9* promoter might impact the expression level of *PhFTL9* and be associated with the climate difference between coastal and inland environments. We further compared nucleotide diversity (π) across the promotor region of *PhFTL9* between coastal ecotypes and central Texas inland ecotypes using resequencing data from 22 *P. hallii* accessions. We observed that π is substantially lower in the inland group than in the coastal group from the start codon to upstream 1 kb promoter regions, while this pattern gradually reversed beyond the upstream 1-1.5 kb promoter region with a precipitous drop in π for the coastal group (Fig. 6C). Finally, phylogenetic footprinting of the promoter region of *FTL9* among orthologs in panicoid grasses revealed two conserved blocks in the upstream region, including 0 - 0.9 kb and 1.4 - 1.8 kb promoter region (Fig. 6D). Interestingly, the 26-bp Indel is located in the non-conserved region in the middle of two conserved blocks (Fig. 6D). This pattern of conservation may point to important regulatory elements of *PhFTL9* that have been maintain by purifying selection in panicoid grasses and potential disruptive polymorphism altering *PhFTL9* regulation.

**Fig. 6.**
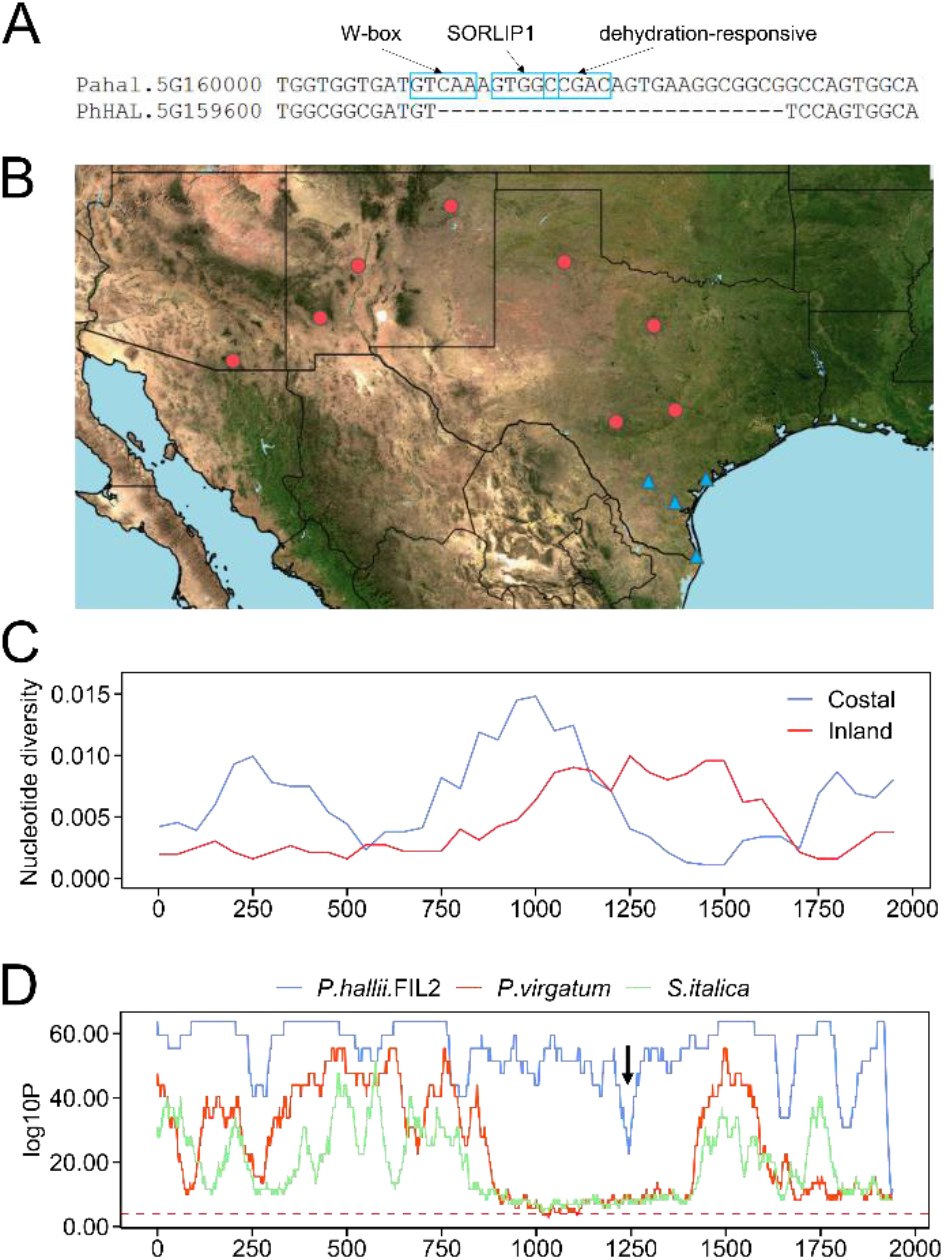
Sequence variation of *PhFTL9* promoter in *P. hallii* and related panicoid grasses. A. A 26-bp InDel in the *PhFTL9* promoter. Blue boxes indicate the *cis*-elements existed in FIL2 allele (Pahal.5G160000). B. The geographic distribution of 12 *P. hallii* accessions. Blue triangles indicate the coastal ecotypes with a 26-bp Indel in the *PhFTL9* promoter, while red circles indicate the inland ecotypes lacking that InDel. C. Sliding-window analysis of nucleotide polymorphism (π) in a sample of inland (red, n=12) and coastal (blue, n=10) of *P. hallii* diversity population. D. Evolutionary conserved regulatory regions of *PhFTL9* homologs in panicoid grasses. The 2-kb promoter sequences from *P. hallii* HAL2 genotype are used as the reference and window size is setting as 60-bp. The *y-axis* gives the log10 transformed *P*-value obtained by aligning one window at this position to any window in *P. hallii* FIL2, switchgrass (*P. virgatum*), and *S*.*italica*. The dashed line indicates the raw score corresponding to the significance threshold for *P* = 0.00001.

## Discussion

### The genetic architecture of flowering time in *P. hallii*

During the past decades, two annual model species, *Arabidopsis* and rice, have served as universal references in studies of the complex network of flowering time genes and how they integrate environmental signals (46, 47). A core regulator involved in the molecular network of flowering is the *Arabidopsis* gene *CONSTANS* (*CO*) and its rice orthologue *HEADING DATE 1* (*Hd1*), encoding zinc-finger transcriptional factors with the *CO, CO-like*, and *TOC1* domains (48, 49). A monocot specific transcriptional activator *EARLY HEADING DATE 1* (*Ehd1*), encoding a B-type response regulator, integrates various molecular signals and induces flowering in diverse environments (50, 51). Both *CO*/*Hd1* and *Ehd1* can bind the promoter regions of genes *FLOWERING LOCUS T* (*FT*) and its orthologues via special *cis*-elements, and regulate the expression of the florigen *FT* to trigger properly timed flowering (50, 52). These advances have facilitated our understanding and insight into mechanisms controlling flowering in *A. thaliana* and in crop domestication.

As a native perennial grass, *P. hallii* germinates and flowers in both spring and fall through the integration of complex seasonal and environmental signals associated with local conditions. Therefore, the genetic basis of flowering time and its potential molecular mechanisms in *P. hallii* could be unique relative to our understanding of flowering time in most annual monocarpic species. Our previous *P. hallii* studies identified two flowering time QTL in an F_2_ population and two panicle emergence QTL in the RIL population with a QTL on chromosome 5 (*qFT-5* in this study) found in both experiments (34, 37). In this study, several major QTL were identified in the F_2_ or RIL population under a given environment, indicating that flowering time variation between HAL2 and FIL2 is impacted by a relatively small number of large effect QTL. This is similar to the results of many native species (e.g. *Arabidopsis*, monkeyflower, and *Brachypodium*), where flowering variation is often impacted by a few loci with major effects (53, 54). Among the major QTL we discovered, *qFT-5* was uniformly detected across different experiments with considerable QTL x E interactions across different environments, signifying an essential role of *qFT-5* in flowering variation in *P. hallii*. Epistatic interaction was detected between the *qFT-5* region and other chromosome regions, suggesting its important role in flowering regulation networks. Combing the results from the fine-mapping and expression analysis, we narrowed the *qFT-5* to 16 annotated genes in a 380-kbp region. There are no other genes known to impact flowering except an orthologue of *FT-like 9* (*FTL9*), suggesting it’s a good candidate underlying *q-FT5*. Although it’s well known that the *FT* gene family is involved in the regulation of flowering time, most studies focus on *FT1* and its orthologues (e.g. *Hd3a* and *RFT1* in rice) as the florigen promoting flowering (39, 41). *FTL9* has not been determined as a cause of natural variation in flowering time in rice, the classic monocot model system. However, this novel *FT* family member has been strongly associated with flowering time in natural populations of switchgrass (55), a close relative of *P. hallii*. Moreover, a recent study suggested that natural variation of *FTL9* conferred competence to flower under short-day vernalization in *Brachypodium* (56). These results suggest a novel role of *FTL9* in flowering variation in temperate grasses.

We also detected a number of small effect QTL that have not been identified before in our earlier mapping studies. Most of these small effect QTL were only identified in specific environments. For example, QTL 1-7/QTL 2-7 and QTL 1-9 are specifically detected under spring or fall short-day conditions. The mapping interval of QTL 1-7/QTL 2-7 and QTL 1-9 spans several important photoperiodic flowering time genes including *Ghd7* and *Ehd1* (23, 50). It is known that *Ehd1* is induced under short-day conditions and functions downstream of *Ghd7* (23). Although we didn’t detect an interaction between QTL 1-7/QTL 2-7 and QTL 1-9, transcriptome data supports the presence of the *Ghd7-Ehd1* regulatory module in *P. hallii* (57). In addition, QTL 3-2.1 and QTL 3-4 are only detected in RIL population in 2016 summer. We observed orthologues of rice *Ghd7*.*1*/*OsPRR37* and *Hd3a* (*FTL2*) located in the mapping interval of QTL 3-2.1 and QTL 3-4, respectively. In rice, *Ghd7*.*1*/*OsPRR37* regulates flowering time and plays an important role in a wide range of adaptation (58, 59), while *Hd3a* is an orthologue of *Arabidopsis FT1* gene, which is a systemically mobile signal to initiate flowering (60, 61). Further fine mapping and studies on the mechanism of these QTL will provide a unique opportunity to understand the genetic architecture of flowering time in perennial temperate grasses.

### Pleiotropy of flowering time genes and its interaction with complex environments

Pleiotropy is defined as one gene affecting multiple traits. This phenomenon is widely observed for genes associated with the flowering time pathway (19). In our study, we observed broad pleiotropic effects of *qFT-5* on growth and fitness-related traits including tiller number, panicle branching number, total flower number and biomass under natural field conditions. Our fine-mapping results and supporting evidence point to *FTL9* as the causal gene underlying these pleiotropic effects. As the integrator of flowering signal networks, it is perhaps unsurprising that *FT* genes often have pleiotropic effects on the coordination of growth and development in addition to flowering. For example, two *FT* family genes, *FT* (*FT1*) and *TSF*, were shown to interact with *BRC1*, a *TB1* clade gene, to modulate lateral shoot outgrowth and repress the floral transition of the axillary buds in *Arabidopsis* (62). Similarly, rice florigen genes, *Hd3a* (*FTL2*) and *RFT1* (*FTL3*), modulates lateral branching and influences yield-related traits under laboratory or agronomic environments (25, 26). These results suggest that the pleiotropy of flowering time genes is broadly evolutionary conserved.

More interestingly, we found that the pleiotropy of *qFT-5* locus is environmentally dependent in our field reciprocal transplant experiment. We only observed pleiotropic phenotypes in the mesic habitat of Kingsville TX but not in the more xeric Austin TX location. A similar environment-dependent pleiotropic effect was reported in the flowering time gene *Ghd7*, which is an upstream regulator of *FT* family genes in rice (63). As a photoperiodic gene, *Ghd7* responses to different environmental stress signals, including ABA and drought (63). Moreover, it has been reported that the *FT* family genes, *Hd3a* and *RFT1*, integrate photoperiodic and drought stress signals to control the flowering time in rice (64). Therefore, one explanation for the environment-dependent pleiotropy of flowering time genes is that these genes differentially respond to complex and often synergistic environmental signals, especially through the integration of signals from light, temperature, and different abiotic stresses. Our recent transcriptome profiling in *P. hallii* has demonstrated that *PhFTL9* expression is photoperiodic (57), but day-length is not the likely driver of the differences we observed between sites given the similar photoperiods of these locations over the course of our experiments. Interestingly, the expression of *PhFTL9* was regulated by a drought treatment in a previous *P. hallii* field experiment (65). Therefore, it is plausible that *PhFTL9* differentially integrates day-length and drought stress signals in *P. hallii* to coordinate flowering time and other development processes at different locations. This type of process may explain the conditional nature of pleiotropy of *qFT-5* as well as its fitness effect. Additionally, research in *Brachypodium* has highlighted the antagonistic role of the *FTL9* orthologue in flowering regulation under different day-length conditions (66). These results suggest that *FTL9* plays a multifaceted role in the fine-tuning and modulation of photoperiodic flowering in temperate grasses. Further study of the molecular function and the dynamic expression of key flowering time genes across different representative natural habitats will improve the understanding of their roles in environment-induced pleiotropy.

### Contribution of flowering time variation in local adaptation

Flowering time often experiences strong selection within and among natural populations and appropriate phenology is widely considered to be critical for successful reproduction and local adaptation (27). Here, we used a test based on segregating trait variance (Fraser 2020) and a reciprocal common garden study to evaluate the impact of selection on flowering time. The observed pattern of QTL effects and the pattern of trait genetic variance in our crosses are consistent with historical directional selection driving the divergence of flowering time between the upland and lowland ecotypes. In contrast, we did not observe a reciprocal home site advantage from our common garden studies, suggesting that the upland and lowland ecotypes did not exhibit fitness tradeoffs during our current study season. While it is common for local genotypes to outperform foreign genotypes in common garden studies (Kawecki and Ebert 2004; Leimu and Fischer 2008), the degree of local adaptation observed is likely to depend on the vagaries of the period of study in ecological time, especially for long lived perennial. For example, we have recently seen evidence of strong local adaptation between upland and lowland ecotypes in this system related to early seedling recruitment (Razzaque and Juenger, in review). It’s possible that fitness related tradeoffs would also be observed at the adult stage if we were to expand to a better estimate of lifetime fitness measured over successive growing seasons or across periods that experience more biotic or abiotic stress. Future field studies with expanded sampling of environments, genetic diversity, and spanning longer ecological time will be valuable complements to our current study of *P. hallii*.

Increasing evidence suggests that the genetic underpinnings of local adaptation are commonly explained by conditionally neutral loci (10). In the case of flowering time, the gain or loss of regulatory motifs may be critical to the accurate sensing of cues and appropriate responses driving the shift from vegetative to reproduction and local adaptation. For example, natural variation of *FT* in *Arabidopsis* is associated with expression differences controlled by variants at *cis*-regulatory sequences, including small sequence polymorphisms (SNP or InDel) or large structural variations (41, 67). In crop plants, a SNP in the promoter of *ZCN8*, a florigen gene in maize, shows a strong association with flowering time and with differential binding by its upstream flowering activator (68). In term of the local adaptation, expression polymorphisms at environmentally sensitive promoter elements may be a common mechanism of conditional neutrality. This hypothesis has been specifically supported in *Arabidopsis* as fitness advantage underlying flowering time mutants is only observed in specific environments. A good example is that drought may modulate the effect of expression variation at the *FRI*, with the functional allele promoting the elevated expression of its downstream gene *FLC* in drier environmental conditions (22, 69).

Here, our study demonstrated conditional neutrality of fitness effects at the *qFT-5* QTL, corresponding to the known biological role of the *FTL9* gene, and likely driven by environmentally sensitive promoter divergence. Interestingly, our previous study reported a trans-by-drought treatment interaction for the abundance of *PhFTL9* transcripts in a drought experiment (65), suggesting that the expression of *PhFTL9* is regulated by a trans QTL interacting with drought stress. In this study, we identified a 26-bp deletion in the promoter of inland alleles of *PhFTL9*, which results in the loss of a number of motifs related to stress responses in inland accessions relative to coastal accessions. Therefore, this polymorphism could be a potential element involved in differential expression regulation across different environments and the cause of the of flowering time divergence. Importantly, our observation of conditional neutrality may be limited to the specific season or environments of our study. How often do flowering time or pleiotropic loci exhibit beneficial, neutral, or deleterious effects? How variable are the fitness impacts of *qFT-5* alleles across spatial variation in habitats, temporal patterns across seasons, or over the span of fitness accrued across the lifetime of a perennial individual? Our study is one of the few showing how individual flowering time QTL contributes to local adaptation in the field. Future studies based on additional field work will helpful clarify the ecological drivers of *qFT-5* fitness effects and will improve our understanding of the molecular underpinnings of adaptation and the evolution of development through pleiotropic effects.

## Materials and Methods

### Plant materials, growth conditions, and phenotyping in greenhouse experiments

*Panicum hallii* F_2_ and RIL population were generated from a cross between two natural inbred accessions HAL2 (*var. hallii*) and FIL2 (*var. filipes*). HAL2 is an accession collected at the Lady Bird Johnson Wildflower Center in Austin, Texas (30.19°N, 97.87°W), which is a seasonally xeric savanna habitat. FIL2 is an accession collected near the Corpus Christi Botanical Gardens in South Texas (27.65°N, 97.40°W), which has mesic coastal prairie habitat. An F_2_ population of 493 individuals was developed for bulked segregant analysis in this study. A subset of the available RIL (212 lines) were employed for genetic mapping studies (37). Seeds of F_2_ and RIL lines were scarified by sandpaper and placed on moistened sand in Petri dishes with a 12 h-light and 12 h-dark photoperiod at 25 °C in the growth chamber. Five days after germination, individual seedlings were transplanted into Promix:Turface:Profile mixture (6:1:1) in square 8.9 cm plastic pots and grown with a randomized design on the bench of the greenhouse at the Brackenridge Field Laboratory at the University of Texas at Austin in Austin, Texas. The germination date of F_2_ population was June 15^th^ (early summer) in 2014 and the average day-length during the F_2_ experiment was 13.7 hours. The germination date of three RIL experiments were September 5^th^ (fall) in 2015, March 3^th^ (spring) in 2016, and July 12^th^ (late summer) in 2016. Flowering time was recorded as the number of days from germination to the day of first panicle emergence from the leaf sheath. Three to eight replicates from each RIL lines were measured for flowering time depending on the space and plant availability. Day-length was calculated as a function of latitude and day of year for the experimental location using formula from Forsythe et al (70). The average day-length during the 2015 fall, 2016 spring, and 2016 late summer RIL experiments was 11.6, 12.6, and 13.3 hours, respectively, obtained from the germination date to the date of the last flowering plant. The day/night temperature was controlled at 28 °C/25 °C in the greenhouse for all experiments.

### Generation and analysis of QTL-seq in F_2_ population

For QTL-seq, two DNA pools, including an early flowering pool (EF-pool) and late flowering pool (LF-pool) were constructed, respectively, by mixing an equal amount of DNA from 100 early flowering (31-38 days to flower) and 100 late flowering (54-68 days to flower) F_2_ plants from the 2014 summer experiment. Pair-end sequencing libraries (read length 150 bp) were prepared by Genomic Sequencing and Analysis Facility in the University of Texas at Austin with an Illumina HiSeq 2500 platform according to the standard manufacturer’s instructions. Quality of the raw reads was assessed using FastQC (http://www.bioinformatics.babraham.ac.uk/projects/fastqc/). Sequencing adapters were trimmed from both pairs of raw reads using Cutadapt (71). The filtered reads from EF-pool and LF-pool were aligned to the HAL2 genome v2.0 assembly using the bwa mem algorithm with default parameters (72). SNP-calling was performed by the mpileup function from bcftools (73). A SNP-index, which is the proportion of SNPs that have different alleles other than the HAL2 reference alleles, was calculated for both EF and LF pool separately following the method described in Takagi et al (74). ΔSNP-index, which is the difference of the proportion of alternative allele between two pools was calculated by subtracting the SNP-index of the EF pool from the SNP-index of the LF pool. The average distributions of the SNP-index and ΔSNP-index for a given genomic interval was estimated by using a sliding window approach with 1 Mb window size and 10 kb step. Confidence intervals were obtained by simulating a F_2_ mapping population with the same pool size of this bulk segregating population with 10000 replications for a given sequence depth. This process was replicated for a range of sequence depths in order to obtain CI precisely for a given sequence depth. The SNP-index graphs for EF-pool and LF-pool, as well as corresponding ΔSNP-index graph were plotted using ggplot2 package in R. Genomic intervals that crossed the 95% CI threshold of ΔSNP-index were considered as candidate genomic regions harboring a locus associated with flowering time.

### Phenotypic evaluation

Pearson correlation was used to estimate the phenotypic correlation of flowering time between seasons. Narrow-sense heritability (*h*^*2*^) was estimated as *V*_*a*_ / *V*_*p*_ using the Sommer package in R, where *V*_*a*_ is the additive variances attributable to genetic relatedness and *V*_*p*_ is the total phenotypic variance (75). *h*^*2*^ was estimated for flowering time in each experiment using the additive kinship matrix, which was obtained based on genotypic information. For genetic correlation estimation, flowering time from three seasonal experiments was used as response variables and similarly the kinship matrix was modeled as a random effect and used to estimate the additive genetic covariance. For G x E interactions on flowering time, we used a likelihood-ratio test to compare the following two models: 1) a main effect model assumed that there is no G x E; and 2) unstructured model assumed that there are G x E interactions and unstructured variance-covariance matrix for the different environments. Significance of the likelihood-ratio test for G x E interactions was assessed at the level of α = 0.05.

### Tests of selection based on segregation in a cross

Fraser (2020) noted that the pattern of trait variance in recombinant populations can provide insight into whether the parental trait divergence is consistent with neutral divergence or more likely the result of natural selection. The approach is a generalization of the QTL sign test (Orr 1998) and rests on the realization that the phenotypic distribution in a segregating/recombinant population can be treated as a null model for the distribution of phenotypes expected under neutral evolution. In this framework, the pattern of underlying QTL effects under strong directional selection is expected to be complementary (in a consistent direction) if the trait has experienced consistent strong directional selection leading to parental divergence, whereas a mixed of positive and negative effects is expected if the trait evolved by neutral processes. By extension, if a trait has experienced strong directional selection driving parental divergence the variance among parental lines is expected to be larger than the segregating variance in the recombinant population, The *v*-test was performed as described in equation 2 of Fraser (2020) for our genetic populations. In brief, to calculate *v*, we first estimated the trait variances within and between parents of the cross. Then we estimated the variance among F_2_ and RIL means and used a *c* value of 2 for F_2_ and 1 for RIL (Fraser 2020).

### QTL analysis in RIL population

The RIL linkage map was built as previously described using the HAL2 genome assembly as the reference. Given our experiments were conducted in different environments, we completed QTL mapping analyses to test for QTL x environment interactions. We compared the likelihood of models allowing QTL to interact with the environment versus reduced models adjusting only for the additive effect of the environment using the R/qtl2 package (76). The full model is expressed as: phenotype = µ + QTL + E + QTL x E + kinship + e, and the reduced model is: phenotype = µ + QTL + E + kinship + e, where µ is the population mean, QTL is the marker effects, E is the environmental effects, QTL x E is the interaction between marker and environmental effects, and e is the error terms. The genome scan was accomplished through ‘scan1’ function. The statistical significance of the genome scan was established by performing a stratified permutation test (n=1000) for the full model and reduced model using ‘scan1perm’ function. The difference of thresholds between these two models from 1000 permutations was considered as the threshold for the QTL x E model. The estimated QTL effect was obtained using ‘scan1coef’ function.

QTL mapping for each RIL experiment was completed in R using a multiple-QTL model in the R/qtl package (77). The ‘scantwo’ function with 1000 permutations was used to calculate penalties for main effect and interactions for each flowering time trait. The ‘stepwiseqtl’ function was used to conduct a forward-backward search with default setting of maximum number of QTL and account for epistasis that optimized the penalized LOD score criterion. Threshold values for type 1 error rates were set at α = 0.05 for flowering time traits were based on permutations. 1.5 LOD drop intervals of QTL were calculated using the ‘qtlStats’ function.

### QTL fine-mapping

We identified a single RIL line (FH-312) that was heterozygous across the 2-LOD support interval of *qFT-5* but homozygous for either HAL2 or FIL2 alleles at the majority of the remainder of the genome. Therefore, this RIL line can be applied to validation of QTL through a heterogeneous inbred family (HIF) strategy (38). We developed a HIF population containing 1,668 F_2_ progenies by self-fertilization of FH-312. Two flanking markers M5-7948 and M5-12422 were used to screen the HIF progenies and identify recombinants. Selfed progeny from selected recombinants were generated to a F_3_ population. Eight additional markers between markers M5-7948 and M5-12422 were used to genotype all recombinant-derived F_3_ individuals. Flowering time of individual plants from the recombinant-derived F_3_ population was scored. Individual marker effects on flowering time were tested using an ANOVA while controlling for initial seedling height. Finally, a pair of NILs (NIL^qFT-5-FIL2^ and NIL^qFT-5-HAL2^) were developed by selfing a heterozygous plant (HIF-2-2-1) and screening with markers to confirm homozygous FIL2 and HAL2 genotypes across the *qFT-5* interval. Primers used for all markers amplification are listed in Table S3.

### Reciprocal transplant field experiments at coastal and inland sites

We used a reciprocal transplant experiment to study QTL allelic effects at the location of origins of the HAL2 and FIL2 parental lines. Our reciprocal transplant experiment was initiated with seedlings. Seeds of HAL2 and FIL2 parental lines and the pair of NILs (NIL^qFT-5-FIL2^ and NIL^qFT-5-HAL2^) created from fine-mapping experiments were scarified by sandpaper and placed on moistened sand in Petri dishes with a 12 h-light and 12 h-dark photoperiod at 25 °C in the growth chamber. Five days after germination, 50 individual seedlings of each genotype were maintained with a randomized design on the bench of the greenhouse at the Brackenridge Field Laboratory at the University of Texas at Austin. Ten days after germination, all seedlings were transplanted to randomized positions in a rectangular array with individual plants separated by 1 m in the field at the UT Brackenridge Field Laboratory in Austin Texas and Kika de la Garza USDA Plant Materials Center in Kingsville Texas on April 5^th^, 2019 and April 7^th^, 2019. We measured a number of traits related to growth and development including tiller number, primary branch number, secondary branch number, number of flowers per panicle, and above-ground biomass at the end of the growing season and associated with fall senescence. We estimated the total growing season fitness from multiplying tillers x number of flowers per panicle. We used a two-way ANOVA to examine the effect of *qFT-5* and site on fitness-related traits and total fitness recorded in the field. To improve normality of residuals and homogeneity of variances, total fitness was square-root-transformed before analysis. When the QTL x site interaction was statistically significant, contrasts were used to examine differences between the local and nonlocal *qFT-5* alleles separately by site.

### DNA extraction and molecular marker analysis

Genomic DNA was isolated using the modified CTAB method from young leaves of F_2_ plants and HIF plants for QTL analysis (78). InDel markers were developed based on HAL2 genome v2.0 and FIL2 genome v3.0 assembly (https://phytozome.jgi.doe.gov/pz/portal.html). Primers were designed using software Primer 3.0 (http://bioinfo.ut.ee/primer3-0.4.0/) and were then amplified from F_2_ individuals and HIF individuals. A total volume of 10 ul reaction mixture was used for PCR amplification, which is composed 1 ul of template DNA, 0.3 ul of each primer (10 mM), 2 ul 5 x Buffer, 1.2 ul MgCl2, 1 ul dNTP, and 0.1 ul GoTag (Promega). Amplification was performed on program for the initial denaturing step with 96 °C for 3 min, followed by 38 cycles for 30 s at 96 °C, 30 s at 58 °C, 30 s at 72°C, with a final extension at 72 °C for 5 min. The PCR products can be well separated using 2.5 % agarose gel electrophoresis.

### RNA extraction and gene expression analysis

The expression level of *PhFTL9* in leaves was measured by quantitative RT-PCR (qPCR). HAL2 and FIL2 individuals were established with a 14 h-light and 10 h-dark photoperiod at 25 °C under greenhouse conditions. Leaf samples were harvested two weeks post germination at 08:00 (zeitgeber time 2) and 18:00 (zeitgeber time 12). We collected the samples from four different plants as three biological replicates for each genotype and stored these samples in liquid nitrogen. For qPCR, we isolated total RNA from the leaves using an RNA extraction kit (TRIzol reagent, Invitrogen). 2 ug total RNA was reverse-transcribed using SSII reverse transcriptase (Invitrogen) in a volume of 80 ul to obtain cDNA. We used primers qPhFTL9-F and qPhFTL9-R for amplifying the transcript of *PhFTL9*. We used primers qPhUBI-F and qPhUBI-R for ubiquitin-conjugating enzyme (Pahal.3G250900) as the internal control. The ubiquitin-conjugating enzyme is a highly expressed gene that was stable across many *P. hallii* tissues (79). We carried out qPCR in a total volume of 10 μl containing 2 μl of the reverse-transcribed product above, 0.25 μM gene-specific primers and 5 μl LightCycler 480 SYBR Green I Master (Roche) on a Roche LightCycler 480 II real-time PCR System according to the manufacturer’s instructions. The relative expression levels of *PhFTL9* were obtained using the delta-delta Ct method (80). Primers used for qPCR analysis are listed in Table S3.

### Sequence analysis

To evaluate protein evolution, we identified the protein-coding sequences of *PhFTL9* orthologues in *Panicum hallii* HAL2 (PhHAL.5G159600), *Panicum hallii* FIL2 (Pahal.5G160000), *Panicum virgatum* (Pavir.5KG539100), *Setaria italica* (Seita.5G317600), *Zea mays* (Zm00001d043461_T001), and *Sorghum bicolor* (Sobic.003G295300) from Phytozome v13 (https://phytozome-next.jgi.doe.gov/). We calculated Ka, Ks, and Ka/Ks values using the software KaKs_Calculator 2.0 (81), with a modified version of the Yang and Nielsen method (82, 83). To explore potential regulatory sequence divergence at *PhFTL9*, we compared the promoter region contained within the upstream 2000 bp from the translation start ATG of *PhFTL9* of the FIL2 v3.0 and HAL2 genome v2.0 assemblies. We aligned these regions using ClustalW2 and characterized known *cis* regulatory elements using the new PLACE database (84). The promoter of *PhFTL9* was amplified using Phusion High-Fidelity DNA Polymerase (New England Biolabs) from genomic DNA. The PCR fragments were purified using PureLink Quick Gel Extraction and PCR Purification Combo kit (Invitrogen) and sequenced by Genomic Sequencing and Analysis Facility in the University of Texas at Austin using Applied Biosystems 3730 DNA Analyzer and BigDye Terminator v3.1 chemistry. Primers used for promoter analysis are listed in Table S3. The EARS (Evolutionary Analysis of Regulatory Sequences), a robust and highly sensitive alignment-based approach, was employed to search for evolutionarily conserved regions in the putative promoter regions of *PhFTL9* and its homologs in *Panicum virgatum, Setaria italica* (85). The promoter region from the HAL2 genome v2.0 assembly was selected as the reference for EARS analysis. To more broadly characterize diversity in the *PhFTL9* promoter we compiled and analyzed resequencing data from a diversity set of natural accessions of *P. hallii* var *hallii* and *P. hallii* var *filipes* collected from natural populations (35). Nucleotide diversity (π) was calculated using a 300 bp window with a step size of 50 bp from SNPs identified from11 inland accessions and 10 coastal accessions using vcftools (86).

## Data Availability

Raw reads for the QTL-seq experiment are available in the Sequence Read Archive database (https://www.ncbi.nlm.nih.gov/sra) with accession number SRR14049075 and SRR14049076. Transcriptome data are available in the Sequence Read Archive database with accession number XXX.

## Acknowledgments

We thank Allison Hutt, Jun Yin, and Maria Jose Gomez for helping with data collection. This research was supported and funded by the National Science Foundation Plant Genome Research Program (IOS-1444533) to T.E.J. The work conducted by the U.S. Department of Energy Joint Genome Institute is supported by the Office of Science of the U.S. Department of Energy under Contract No DE-AC02-05CH11231.

## Summary of supplementary information

**Fig. S1**. Genotypic effects of the five QTL with QTL x E detected in the full model. AA and BB indicate FIL2 and HAL2 alleles, respectively. These allelic effects are plotted side-by-side for each seasonal experiment. Positive additive effects indicate a delayed flowering time from that allele, while negative effects indicate an accelerated flowering time from that allele.

**Fig. S2**. Pairwise epistatic QTL in the 2016 summer RIL population. Plotted points indicate two-locus genotype means ± SE for the two loci containing flowering time between QTL-3-5 and QTL-3-1. AA and BB indicate homozygous FIL2 and HAL2 genotypes at relative QTL loci, respectively.

**Fig. S3**. Graphical genotypes of a heterogeneous inbred family (HIF) (FH-312) used for used for constructing Near Isogenic Lines (NILs) for fine mapping of *qFT-5*. Blue indicates regions homozygous for FIL2; red indicates regions homozygous for HAL2; green indicates heterozygous regions; grey indicates unknown regions. The *qFT-5* region are indicated by black circle.

**Fig. S4**. The flowering time comparison between HIF progenies carrying FIL2 homozygous alleles at *qFT-5* region than those carrying HAL2 homozygous alleles. AA indicates the HIF progenies carrying homozygous FIL2 genotypes at two flanking markers (M5-7948 and M5-12422), while BB indicates the HIF progenies carrying homozygous HAL2 genotypes at two flanking markers (M5-7948 and M5-12422). The values are shown in Mean (SE) with n > 200.

**Fig. S5**. Expression analysis of *PhFTL9* gene between HAL2 and FIL2. Leaf samples were harvested two weeks post germination at 08:00 (zeitgeber time 2, cornsilk bars) and 18:00 (zeitgeber time 12, gray bars). Bars and error bars indicate mean values and SE, respectively, based on four biological repeats.

**Fig. S6**. The alignment of PhFTL9 protein sequences between FIL2 (Pahal.5G160000) and HAL2 (PhHAL.5G159600) alleles.

**Fig. S7**. The alignment of *PhFTL9* promoter sequences (∼2kb) between FIL2 (Pahal.5G160000) and HAL2 (PhHAL.5G159600) alleles. The 26-bp Indel is indicated by red rectangle.

**Table S1**. Performance of flowering time in parents, F_2_, and RIL population.

**Table S2**. Directional selection of flowering time divergence using Fraser v-test.

**Table S3**. Primers used in this study.

**Table S4**. QTL-by-environment interaction in RIL population.

**Table S5**. Main effects of QTL at each RIL experiment.

**Table S6**. The list of annotated genes in 380-kb interval of NIL plants.

**Table S7**. ANOVA test in reciprocal transplant experiment.

**Table S8**. Summary of Ka, Ks, and Ka/Ks of *PhFTL9* orthologous pairs in panicoid grasses.

## Notes

### Competing Interest Statement

The authors have declared no competing interest.

